# Prior cocaine exposure increases firing to immediate reward while attenuating cue and context signals related to reward value in anterior insula

**DOI:** 10.1101/2020.10.01.322776

**Authors:** Heather J. Pribut, Daniela Vázquez, Adam T. Brockett, Alice D. Wei, Stephen S. Tennyson, Matthew R. Roesch

## Abstract

The insula contributes to behavioral control and is disrupted by substance abuse, yet we know little about the neural signals underlying these functions or how they are disrupted after chronic drug self-administration. Here, rats self-administered either cocaine (experimental group) or sucrose (control) for twelve consecutive days. After a one-month withdrawal period, we recorded from anterior insula while rats performed a previously learned reward-guided decision-making task. Cocaine-exposed rats were more sensitive to value manipulations and were faster to respond. These behavioral changes were accompanied by elevated counts of neurons in the insula that increased firing to reward. These neurons also fired more strongly at the start of long delay trials-when a more immediate reward would be expected, and fired less strongly in anticipation of the actual delivery of delayed rewards. Although reward-related firing to immediate reward was enhanced after cocaine self-administration, reward-predicting cue and context signals were attenuated.

**Significance Statement:** The insula plays a clear role in drug-addiction and drug-induced impairments of decision-making, yet there is little understanding of its underlying neural signals. We found that chronic cocaine self-administration reduces cue and context-encoding in insula, while enhancing signals related to immediate reward. These changes in neural activity likely contribute to impaired decision-making and impulsivity observed after drug use.

## INTRODUCTION

The insula has recently gained traction as a key contributor to relapse and drug-seeking behaviors, and as a potential therapeutic target for addiction. Work in humans has shown that insula is activated during the presentation of drug cues and recollection of past drug use (Bonson et al., 2002; Wang et al., 1999). Further, it has been shown that damage to insula can promote drug abstinence (Naqvi et al., 2007). Rodent work has supported these ideas by showing that disruption of insula function reduces the ability of drug-associated cues and contexts to drive drug-seeking (Campbell et al., 2019; Cofresí et al., 2018; Contreras et al., 2007; Forget et al., 2010; García et al., 2013; Gil-Lievana et al., 2020; Li et al., 2013; Pelloux et al., 2013; Perdomo et al., 2012; Pushparaj and Le Foll, 2015; Seif et al., 2013; Zhang et al., 2019).

Although it is clear that insula is directly involved in drug addiction, it is less clear how it contributes to normal behavioral control, or how those mechanisms might be disrupted by chronic drug abuse. Traditionally, the insula has been described as an interoceptive center (Craig, 2010, 2002; Damasio et al., 2000; Naqvi and Bechara, 2010), but is also thought to contribute to functions related to reward processing and decision-making (Burke and Tobler, 2011; Droutman et al., 2015; Mesulam and Mufson, 1982; Ongur and Price, 2000; Preuschoff et al., 2008; Rogers-Carter and Christianson, 2019). Consistent with these functions, recent work has shown that firing in insula correlates to the anticipation and delivery of both positive and negative outcomes (Guillem et al., 2010; Jo and Jung, 2016; Kusumoto-Yoshida et al., 2015; Mizoguchi et al., 2015; Moschak et al., 2018; Samuelsen et al., 2012; Vincis et al., 2020; Wittmann et al., 2020). Here, we ask if outcome-related neural correlates in insula are disrupted by chronic cocaine-self administration in rats performing a reward-guided decision-making task.

During performance of this task (*Fig. 1A,B*) rats use reward-predicting cues to guide choice behavior between two options, the values of which change regularly. The values of the two options are manipulated across two contexts: in one context, rats choose between an immediate and delayed reward; in the other context, rats choose between a large and small reward with delays to reward held constant. In both contexts, rats learn which option yields the preferred outcome, and maintain those expectations during delays to reward and across trial blocks while simultaneously following forced-choice rules. We have shown across several studies that rats prefer immediate over delayed reward, and large over small reward, and that rats are overall more motivated during the ‘size’ block context (Burton et al., 2018b, 2017; Roesch et al., 2007; Roesch and Bryden, 2011; Vázquez et al., 2019). Further, we have shown that rats that had previously self-administered cocaine for two weeks are more sensitive to value manipulations and exhibit faster reaction times months after drug exposure (Brockett et al., 2018; Burton et al., 2018a, 2017; Vázquez et al., 2019).

**Figure 1.**
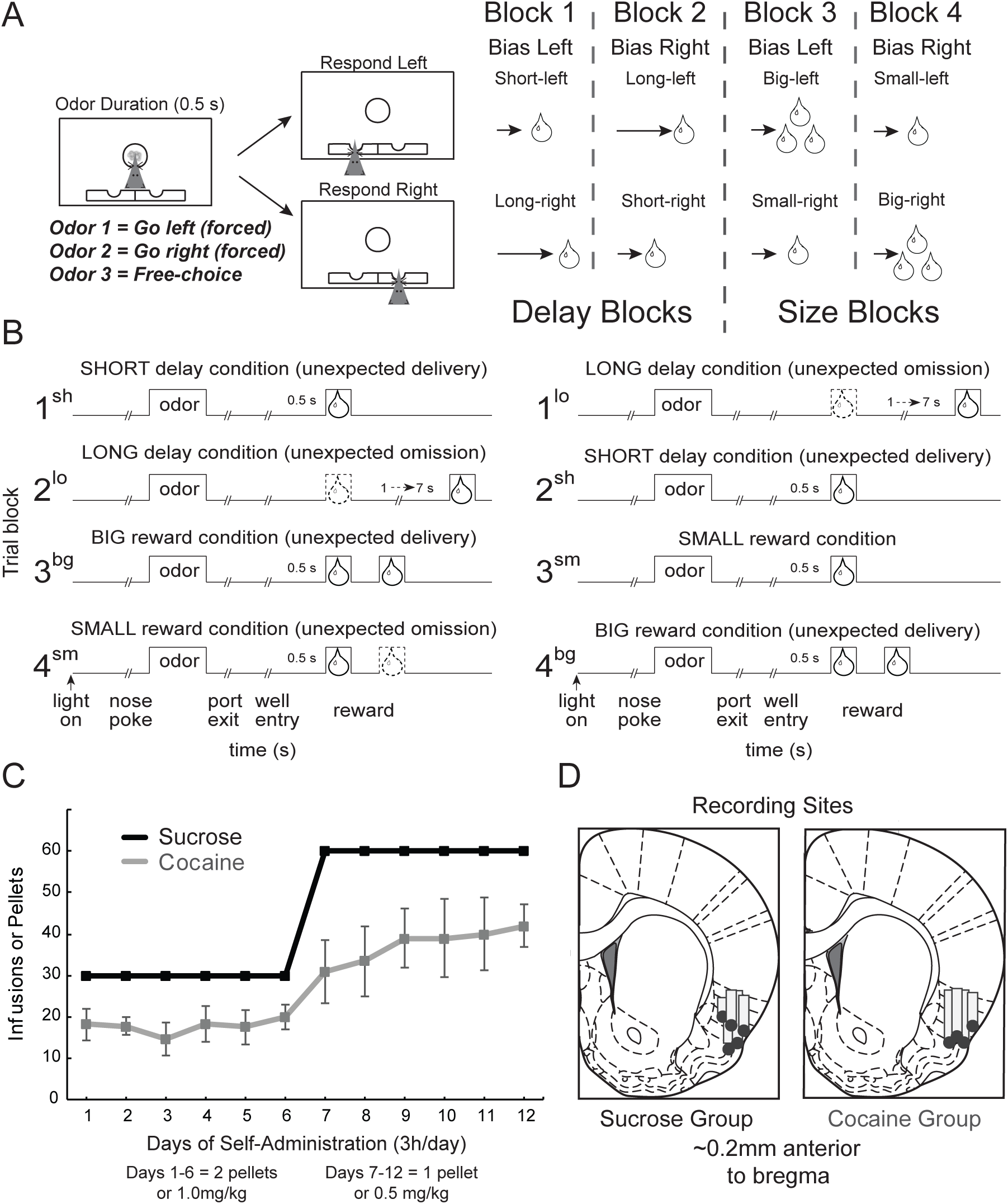
Reward-guided decision-making task, self-administration data, and recording sites. **A**. Task schematic, showing an example trial (left panels) and block sequence in a session (right). Rats nose-poked in a central port for 0.5s to receive an odor with a duration of 0.5s. Odors would instruct them to respond to fluid wells below where they could receive liquid sucrose rewards after 500-7000ms. **B**. Summary of block structure. During a recording session, one fluid well would be arbitrarily designated as short (delivering reward after a short 500ms delay) and the other designated as long (delivering reward after a 1-7s delay). Contingencies were reversed after ∼60 trials (Block 2). In Block 3, delays to reward were held constant at 500ms, but the well designated as long in the previous block now offered 2 boli (i.e. big reward), while the other well offered only 1 (i.e. small reward). Contingencies switched once again in Block 4. Each block shift was unsignaled so that rats needed to learn through behavior that reward values had changed. **C**. Average lever-press rates for Control (black) and Cocaine-exposed (gray) rats for each day of self-administration. Data points represent each day. Bars on the cocaine-exposed group represent standard error of the mean (SEM). **D**. Electrode placement for each rat, verified by histology.

Given insula’s role in addiction and its dysregulation following drug use, we hypothesized that functional signals related to reward processing would be disrupted by chronic cocaine self-administration. We found that reward delivery-related signals in cocaine-exposed rats were enhanced, while reward context- and cue-related firing was reduced. Further, encoding of more immediate delivery of reward was more prominent in cocaine-exposed rats. Overall, these results suggest that heightened reward responding and lower cue and context selectivity in insula may contribute to increased impulsivity and poor decision-making observed after chronic drug use (Brockett et al., 2018; Burton et al., 2018b; Mendez et al., 2010; Roesch et al., 2007; Simon et al., 2007; Vázquez et al., 2019).

## RESULTS

### Self-Administration

Lever-press activity for the active lever was recorded across all 12 days of self-administration for each rat (*Fig. 1C*). During weeks 1 and 2, rats in the cocaine group self-administered an average of 17.75 (standard deviation = 8.61) and 37.42 (standard deviation = 18.88) infusions, respectively. Control rats that pressed for sugar pellets maxed out during every session. Both cocaine and control rats responded at significantly higher rates during the second week of self-administration (*ttest*, t(23) = 5.507, *p* < 0.001).

### Cocaine-exposure produced a response-bias towards high-value rewards

Behavioral data from 416 cocaine recording sessions (n’s for each rat = 84, 128, 115, 89) and 315 control sessions (n’s = 75, 81, 74, 72, 13; Fig. 1D) were analyzed using a multi-factor ANOVA across dependent variables of percent choice, percent correct, and reaction times on forced- and free-choice trials. The ANOVA factors consisted of group (control vs cocaine), value (high vs low), and block (delay vs size).

Analysis of free-choice trials (*Fig. 2A*) found a significant main effect of value (F(1,5841) = 1077.58, *p* < 0.001), demonstrating a greater preference for high-valued rewards across blocks in both cocaine and control rats; selection of high value rewards was significantly greater in both delay (*ttest*, t(1461) = −53.458, *p* < 0.001) and size blocks (*ttest*, t(1461) = −21.551, *p* < 0.001). Cocaine rats’ stronger preference for high value rewards is further illustrated in the inset showing the difference between high (i.e., short; big) and low (i.e., long; small) value free-choice responding for control and cocaine-exposed rats (*Fig. 2A*; inset; *ttest*, t(707) = −5.026, *p* < 0.001).

**Figure 2.**
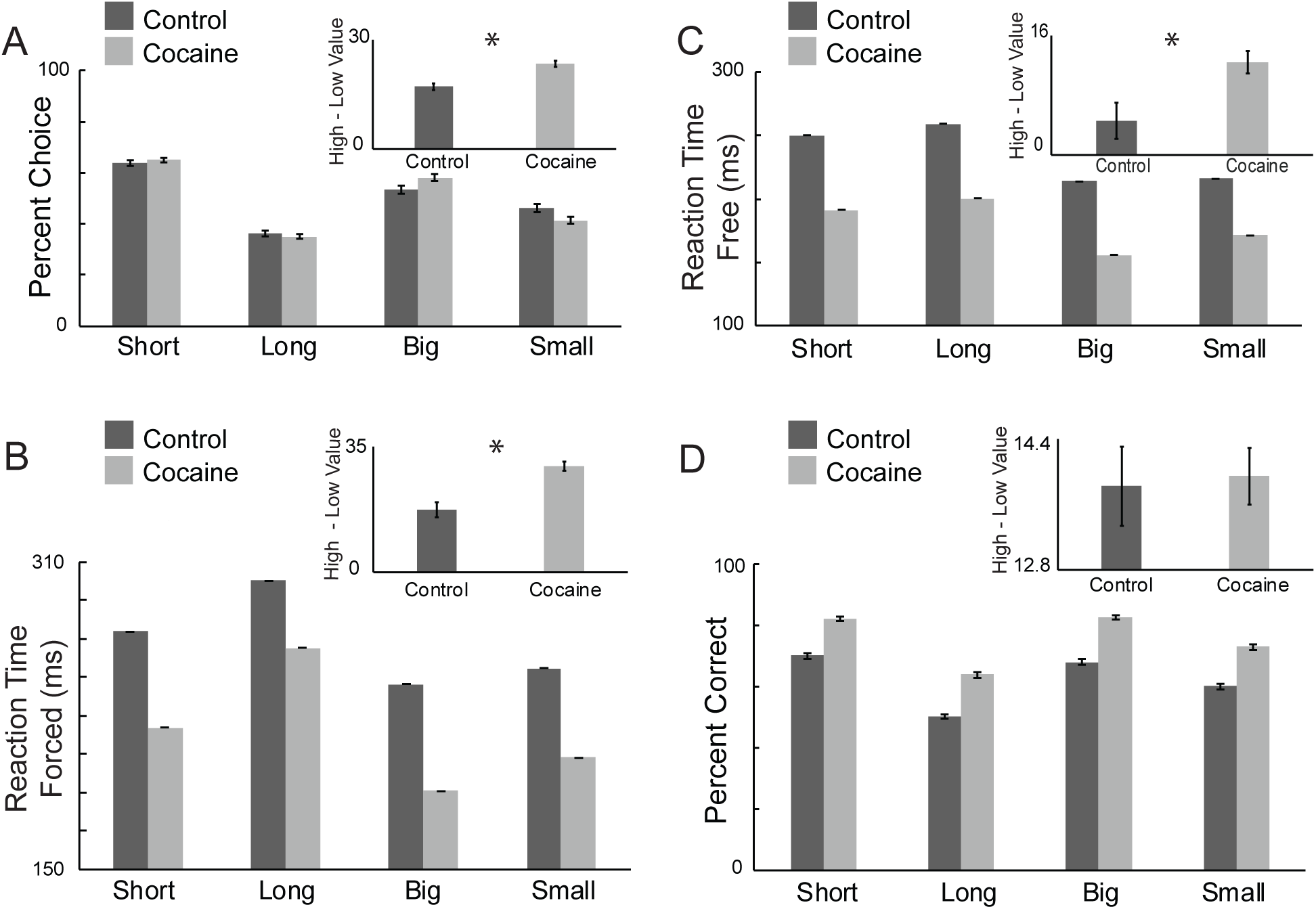
Cocaine exposure produced faster reaction times and response biases for high-valued rewards. All error bars indicate SEM. **A**. Percent choice on all free-choice trials in each value manipulation. Results were averaged across animals and weighted by the number of recordings performed to provide a better representation of the behaviors observed as related to the neural analysis (controls, dark bars; cocaine, light bars). Inset: Response bias (percent choice high – low value rewards). **B**. Reaction time (odor port exit – odor onset) on forced-choice trials for each value manipulation. Inset: Response bias (reaction time low – high value rewards) **C**. Reaction time for free-choice trials. Inset: Response bias, calculated as in **C. D**. Percent correct on all forced-choice trials. Inset: Response bias, calculated as in **A**.

This cocaine-induced exaggerated response bias toward higher valued reward was also observed in reaction times. Main effects of value and block during forced-choice trials demonstrated that all rats were faster for high-valued rewards (*Fig. 2B*; F(1,5841) = 69.26, *p* < 0.001), and were also faster for size blocks compared to delay blocks (F(1,5841) = 209.49, *p* <0.001). There was also a main effect of cocaine indicating that rats that had self-administered cocaine were altogether faster compared to controls (F(1,5841) = 276.87, *p* < 0.001). Finally, cocaine-exposed rats showed a stronger response bias, as illustrated in the inset that plots the difference between reaction times for high and low value rewards (*Fig. 2B*, inset; significant interaction between group and value F(1,5841) = 4.7, *p* = 0.030); *ttest*, t(543) = 4.856, *p* < 0.001). Thus, cocaine-exposed rats exhibited an exaggerated response bias toward high-value rewards, as well as faster reaction times.

Similarly, reaction times during free-choice trials (*Fig. 2C*) were faster for high-valued rewards (Main effect of value; F(1,5717) =11.05, *p* < 0.001) and during size blocks compared to delay blocks (Main effect of block; F(1,5717) = 186.14, *p* < 0.001). Rats that had self-administered cocaine were also significantly faster compared to controls overall (Main effect of group: F(1,5841) = 443.5, *p* < 0.001), particularly for high-valued rewards (*Fig. 2C*; inset; *ttest*, t(537) = 2.733, *p* = 0.006).

Altogether, these results replicate previous findings that rats demonstrate a preference for higher-valued rewards through their choices and reaction times, and that responding is biased towards high-valued rewards in animals previously exposed to cocaine (Brockett et al., 2018; Burton et al., 2018b, 2017; Vázquez et al., 2019). Interestingly, in contrast to previous studies, we found that cocaine-exposed rats were overall more accurate on forced-choice trials (*Fig*.*2D*). There were significant main effects of block and value, with rats being more accurate during size blocks (F(1,5841) = 90.09, *p* < 0.001) and for high-valued rewards (F(1,5841) = 945.27, *p* < 0.001). An additional main effect of group showed that cocaine rats altogether followed forced-choice rules more accurately compared to controls (F(1,5841) = 866.17, *p* < 0.001), more strongly adhering to stimulus-response (S-R) contingencies (i.e., odor 1 = go left; odor 2 = go right).

### Cue-related value-encoding was attenuated after cocaine exposure

In the first analysis, we asked if cocaine-exposure altered the counts of insula neurons that were responsive during odor sampling (odor epoch: 100 ms after odor onset to port exit) compared to baseline (1s before odor onset; *Wilcoxon*, p < 0.05). In controls, 28% (n = 88) of neurons were responsive during odor sampling (*Fig. 3A*), whereas only 12% (n =51) were responsive in rats that had self-administered cocaine (*Fig. 3B*). Both counts were significantly greater than expected by chance alone (control: *chi-square* = 617.371, *p* < 0.001; cocaine: *chi-square* = 135.51, *p* < 0.001), but the frequency of cells that increased firing in controls was significantly higher compared to cells from cocaine-exposed rats (*chi-square* = 18.395, *p* < 0.001). Thus, rats that had self-administered cocaine had fewer cue-responsive neurons compared to controls. An example of a cue-responsive neuron is illustrated in *Figure 3C*. Many of these neurons were value selective, firing more strongly for odor cues that predicted a larger reward or shorter delay for behavioral responses to be made into the cells response field (i.e., the cell’s preferred response direction, or the direction that elicited the strongest firing; e.g., left panels in Figure 3C).

**Figure 3.**
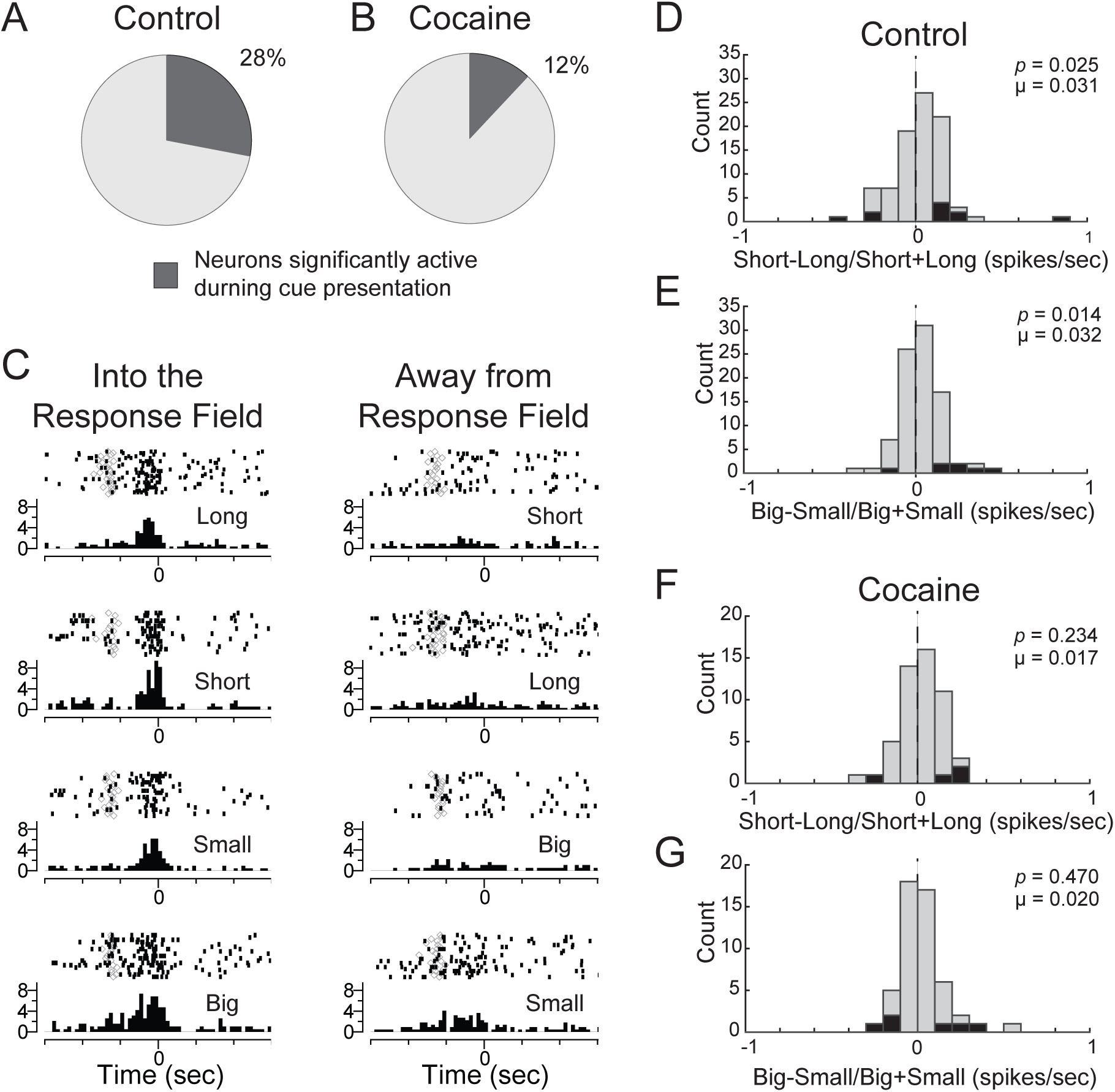
Cocaine exposure reduced cue-induced activity and value-encoding from odor-response cells. Selective vs non-selective odor-responsive cells from **A**. control and **B**. cocaine-exposed rats. Selectivity was defined by significantly greater firing rates during the odor analysis epoch (100ms after odor onset to odor port exit) compared to the baseline analysis epoch (1s before odor onset; *Wilcoxon, p* < 0.05)). **C**. Raster plots from an odor-responsive example cell. Activity is aligned to odor onset, with each row displaying activity for each reward-type. **D**. Distribution of value analysis (short – long/ short + long; big – small/ big + small) for cells taken from control rats, comparing firing rates for short and long-delay rewards. **E**. Similar analysis as **D**, comparing firing rates for big and small rewards during blocks where reward size is manipulated. **F**,**G**. Same analyses as **D**,**E**, but for cells from cocaine-exposed rats.

To quantify value selectivity during the odor epoch, we computed the difference between firing for high and low-value rewards in each neuron during this epoch (i.e., odor onset to port exit) for delay (delay index = short – long / short +long) and size blocks (size index = big – small / big + small), for responses made into each cell’s response field (i.e., the direction the elicited the strongest response). For controls, there were significant positive shifts in both delay (*Wilcoxon, p* = 0.025, µ = 0.031; *Fig. 3D*) and size (*Wilcoxon, p* = 0.014, µ = 0.032; *Fig. 3E*) indices, indicating a higher frequency of neurons with stronger firing rates to cues that predicted high-value reward within both size and delay trial blocks. In rats that had self-administered cocaine, there were no significant shifts in either delay (*Wilcoxon, p* = 0.234, µ = 0.017, *Fig. 3F*) or size indices (*Wilcoxon, p* = 0.470, µ = 0.020, *Fig. 3G*). Although neither delay nor size index cocaine distributions were significantly shifted, neither were significantly different from control distributions (*Wilcoxons*; delay: z = −0.754, *p* = 0.451; size: z = −0.889, *p* = 0.374). Thus, we conclude that cocaine self-administration diminished value-encoding during odor sampling by reducing the counts of odor-responsive neurons (*Fig. 3A* vs *B*), and only slightly attenuating outcome selectivity in those that remained (*Fig. 3D, E* vs *Fig. 3F, G*).

### Immediate rewards were more strongly represented after cocaine exposure

Our next analyses examined neurons whose activity increased during the anticipation and delivery of reward. In controls, we found 46% (n = 144) of neurons increased firing during the reward epoch (*Fig. 4C*; reward epoch: 250ms before reward delivery to 1s after reward delivery) compared to baseline (*Wilcoxon, p* < 0.05. *Chi-square greater than chance* = 1568.594, *p* < 0.001). In rats that had self-administered cocaine, we found an 18% increase in the counts of reward-responsive neurons (*Fig. 4D*; 64%; n = 265). The proportion of reward-responsive neurons in cocaine-exposed rats was significantly greater than the proportion observed in controls (*chi-square* = 6.454, *p* = 0.011).

**Figure 4.**
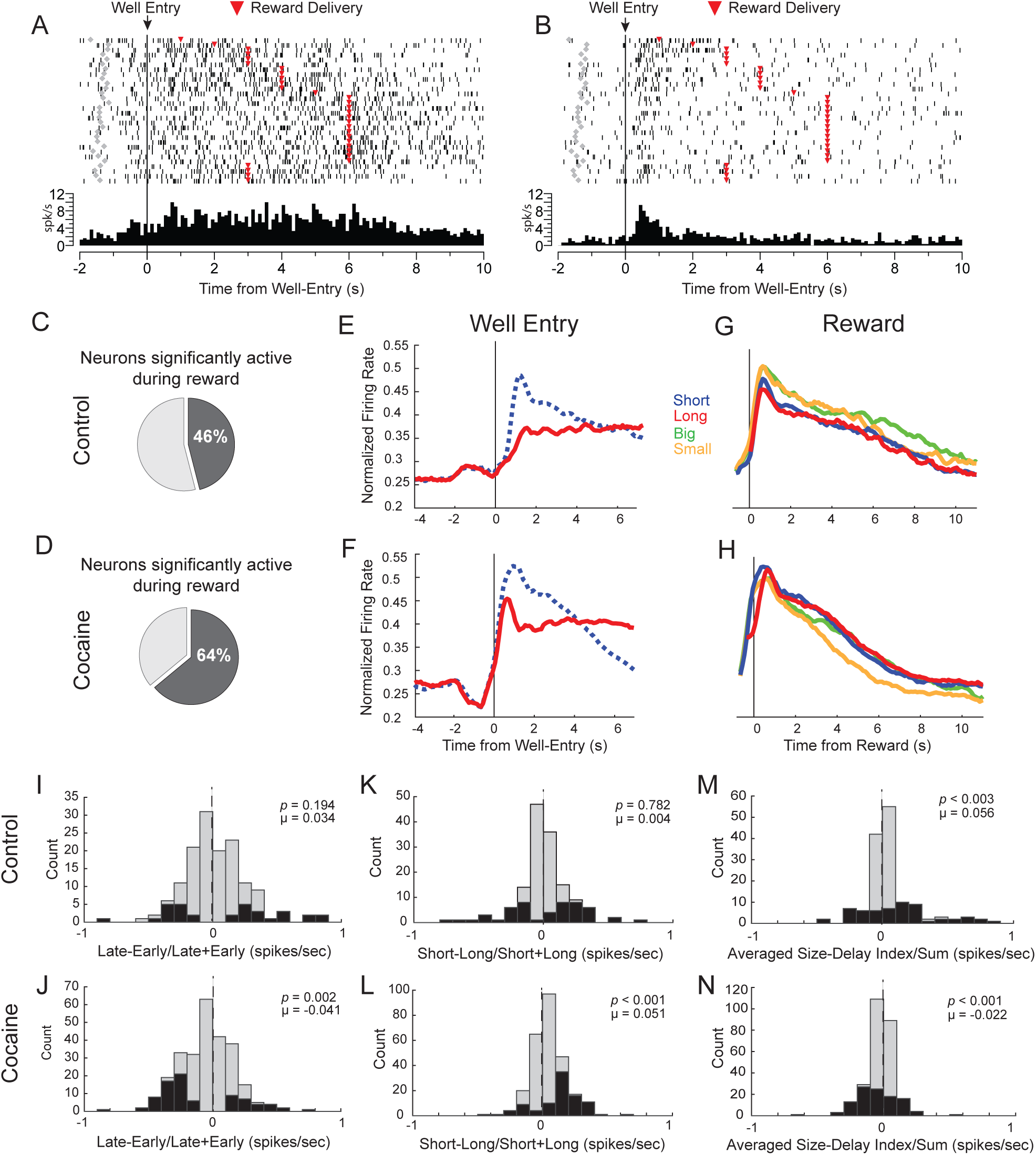
Cocaine exposure reduced delay-representation and activity was discounted for long-delay rewards in reward-responsive cells. **A**. Single neuron example during a long-delay trial in which activity is sustained during the delay period. Activity is aligned to well-entry (time 0, black line), **B**. Single neuron example during a long-delay trial in which activity initially increases after well-entry, and subsequently decreases, failing to maintain reward-representation throughout the long delay. **C**. Percentage of reward-selective cells from control and **D**. cocaine-exposed rats, as defined by significantly greater firing during the reward analysis epoch (250ms before reward to 1s after reward delivery) compared to baseline (1s before odor onset; *Wilcoxon, p* < 0.05). **E**. Normalized firing rates for neurons that increased firing rates during the reward epoch from control (n = 144) and **F**. cocaine-exposed rats (n = 265). Activity is aligned to fluid well entry, comparing short (blue dotted line) and long-delay (red line) rewards. **G**. Normalized firing rates for cells from control and **H**. cocaine-exposed rats, aligning activity for short (blue), long (red), big (green), and small (orange) rewards to reward delivery. **I**. Distribution of value indices for cells taken from control and **J**. cocaine-exposed rats, comparing firing rates during the first and last 500ms of long delay rewards. **K**. Distribution of delay indices (short – long/ short + long) for cells taken from control and **L**. cocaine-exposed rats, comparing firing rates during the reward epoch. **M**. Distribution of context indices reflecting changes in global value (size block – delay block/ size block + delay block) comparing average activity during size and delay blocks during the post reward delivery epoch (1-2 s after reward delivery), for control and **N**. cocaine-exposed rats.

Single cell examples from this population of neurons are illustrated in *Figure 4A* and *B*. As shown in *Figure 4A*, many neurons fired in anticipation of reward across long delays, exhibiting sustained firing from well entry until reward delivery. However, other neurons, as illustrated in *Figure 4B*, seemed to fail in representing reward across a long delay. These cells exhibited increases in firing upon well entry, at the time when reward would have been delivered on the majority of trials (i.e., after 500 ms), followed by a decrease in firing during the remainder of the delay. To determine the response patterns across the entire population of reward responsive neurons, we plotted average firing aligned to both well entry (*Fig. 4E,F*) and reward delivery (*Fig. 4G,H*) for actions made into the response field.

Aligning activity to the well entry, we found that insula neurons from control rats exhibited sustained elevated firing during long delays starting at well entry (*Fig. 4E*; red). Notably, in controls, anticipatory firing after well entry for longer delays (red) rose less rapidly than for rewards delivered after a short delay (blue dashed). This was not true after cocaine exposure (*Fig. 4F*). Instead, early activity during long delays tracked firing as if on a short delay trial, peaking around the time when the more immediate reward would have been delivered before dropping to sustained levels. Thus qualitatively, it appeared that firing after cocaine-exposure held on to the expectation that reward was going to be delivered after a shorter delay (which occurs on 3 (short; big; small) of the 4 trials-types).

To quantify this observation, we computed the difference in activity during the first and last 500ms of the delay period on long-delay trials (Late – Early / Late + Early; early = first 500ms after well-entry; late = last 500ms before reward). These two time points encompass non-overlapping activity early and late in the delay period, when rats were maintaining hold in the fluid well before reward delivery. In controls, we observed no significant shift in the distribution, demonstrating that both early and late expectancy-related firing were similarly represented across longer delays (*Wilcoxon, p* = 0.194, µ = 0.034, *Fig. 4I*). However, for rats that had self-administered cocaine, this distribution was significantly shifted below zero (*Wilcoxon, p* = 0.002, µ = 0.041, *Fig. 4J*) and significantly different from the control distribution (*Wilcoxon*, z = −2.866, *p* = 0.004), indicating that cocaine-exposed rats had a higher frequency of neurons firing more strongly at the start compared to the end of long delays.

From these results it appears that in cocaine-exposed rats, neural correlates of immediate reward expectancy were maintained during performance of long-delay trials. Our findings also suggest that cocaine exposure impairs the ability of insula neurons to shift expectancy-related firing from early to late in delay trials. Such a lack of firing might lead to a reduced or discounted neural representation of delayed rewards compared to those delivered after 500ms (i.e., short-delay trials). Overall, this result would suggest that more immediate rewards are better represented in insula after cocaine exposure. Indeed, examination of average firing aligned to reward delivery (*Fig. 4G,H*) suggests that anticipatory firing for immediate rewards (i.e., those delivered after short delays; blue) was stronger compared to firing for delayed rewards (*Fig. 4H*, red). To quantify this effect, we computed the delay (short-long / short+long) indices for each neuron during the reward epoch. The distribution of delay indices (short-long / short+long) from rats that had self-administered cocaine were significantly shifted in the positive direction (*Wilcoxon, p* < 0.001, µ = 0.051, *Fig. 4L*) and significantly different from the control distribution (*Fig. 4K, Wilcoxon, p* = 0.782, µ = 0.004; *Wilcoxon*, z = 3.081, *p* = 0.002).

Together, these results suggest that previous cocaine exposure increased neural signals for immediate rewards during shorter delay trials, but also over-represented the anticipation of more immediate reward, both within long delay trials themselves and when directly comparing immediate to delayed reward at the time of delivery.

### Context-related value signals were attenuated by cocaine exposure

Upon constructing the population histograms for the analysis above (*Fig. 4G,H*), we unexpectedly found that neurons in insula appeared to encode block context in control rats. Specifically, firing was higher after reward delivery during size blocks (green and orange vs blue and red). Remarkably this was true even for ‘small’ rewards (orange) that were physically the same delay and size as reward on ‘short’ delay trials (blue).

To quantify this effect, we created a ‘context’ index comparing average activity from size and delay blocks (Size Block – Delay Block / Size Block + Delay Block) 1-2s after reward that indexed the global value differences between blocks (*Fig*.*4M,N*). This epoch of interest occurs after the completion of the trial, while rats are still in the well consuming reward but are aware of the reward that was delivered on that trial. From this analysis we found insula neurons exhibited a positive shift in distributions of activity, confirming this unexpected novel finding of higher firing after reward delivery during size compared to delay blocks at the level of single neurons (*Wilcoxon, p* = 0.003, µ = 0.056; *Fig. 4M*). Interestingly, this effect was not observed in cocaine-exposed rats. Instead, the context index distribution was significantly shifted below zero, indicating that cells tended to fire more strongly for delay blocks compared to size blocks (*Fig. 4N*; *Wilcoxon, p* = 0.001, µ = −0.022. Cocaine vs control: *Wilcoxon*, z = −4.287, *p* < 0.001).

Remarkably, we also observed increases in firing during size compared to delay trial blocks in cells that fired more strongly prior to the trial’s start. This finding is illustrated in *Figure 5A*, which plots the average firing of 69 control neurons (22% of total). These neurons ramped up firing in anticipation of house light onset (i.e., the stimulus that signaled the start of each trial) and fired more strongly during ‘size’ compared ‘delay’ trial blocks. To quantify this effect, we again computed the ‘context’ index (Size – Delay / Size + Delay) during the 4s prior to light onset (the period of time immediately preceding the onset of the trial; *Fig. 5B*). Consistent with observations in the population histogram, we found a significant positive shift in distributions, demonstrating that insula neurons tended to fire more strongly during size compared to delay blocks prior to trial onset (*Wilcoxon, p* < 0.001, µ = 0.062).

**Figure 5.**
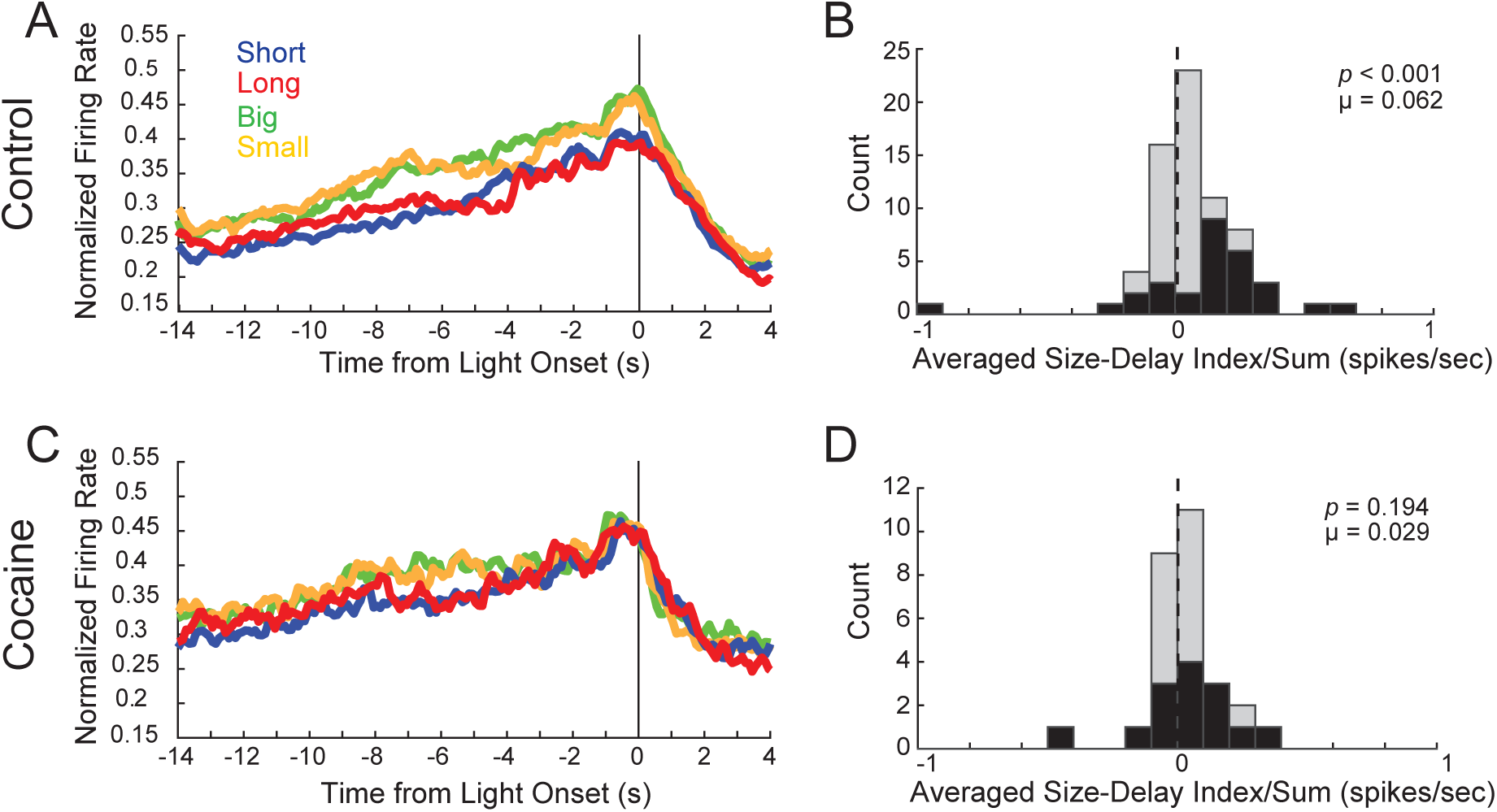
Cocaine-exposure reduced context encoding. **A**. Normalized firing rates for cells from control rats that exhibited elevated activity prior to trial onset. **B**. Distribution of context indices for controls (n = 69) reflecting changes in global value (size block – delay block/ size block + delay block) comparing average activity during size and delay blocks during the pre-trial epoch (4 s before light onset). **C, D**. Same as A and B for cocaine-exposed rats (n = 28).

Interestingly, cells that carried this signal were nearly abolished in rats that had self-administered cocaine (*Fig. 5C*). Only 7% (n = 28) of neurons exhibited such pre-trial ramping activity, which were significantly fewer cells compared to such neurons found in controls (*chi-square* = 25.963, *p* < 0.001). For those cells, the Size-Delay index distribution was not significantly shifted (*Wilcoxon, p* = 0.194, µ = 0.029, *Fig. 5D*), and was not significantly different compared to controls (*Wilcoxon*, z = −1.071, *p* = 0.284). Thus, we conclude that cocaine self-administration diminished contextual pre-trial value-encoding by reducing the counts of responsive neurons and slightly attenuating selectivity in those that remained.

### Contextual block signals correlated with changes in behavior in control rats

Notably, we have not observed these context signals in any other brain area that we have recorded from in rats performing this task (Brockett et al., 2018; Bryden et al., 2011; Burton et al., 2018b, 2017, 2015,2014b, 2014a; Kashtelyan et al., 2012; Roesch et al., 2007, 2006, 2012; Roesch and Bryden, 2011; Takahashi et al., 2017; Vázquez et al., 2019). Thus, context-related signals related to delay and size blocks appear to be unique to insula. To better understand the relationship between context-related firing and behavior, we correlated differences in firing between ‘size’ and ‘delay’ blocks to differences in percent corrects and reaction times between ‘size’ and ‘delay’ blocks (Size – Delay / Size + Delay). Recall that rats were significantly faster and better on ‘size’ compared to ‘delay’ blocks.

In control rats, there was a significant positive relationship between size vs delay block accuracy and size-encoding (*Regression, p* < 0.001, *r*^2^ = 0.095;). Thus, increases in firing were correlated with better performance. Interestingly, after cocaine exposure there was no significant correlation between firing and percent correct (*Regression, p* = 0.808, *r*^2^ < 0.001; *Fig. 6A*; *Fisher r-to-z transformation* comparing control to cocaine; z = 2.9, *p* = 0.002).

**Figure 6.**
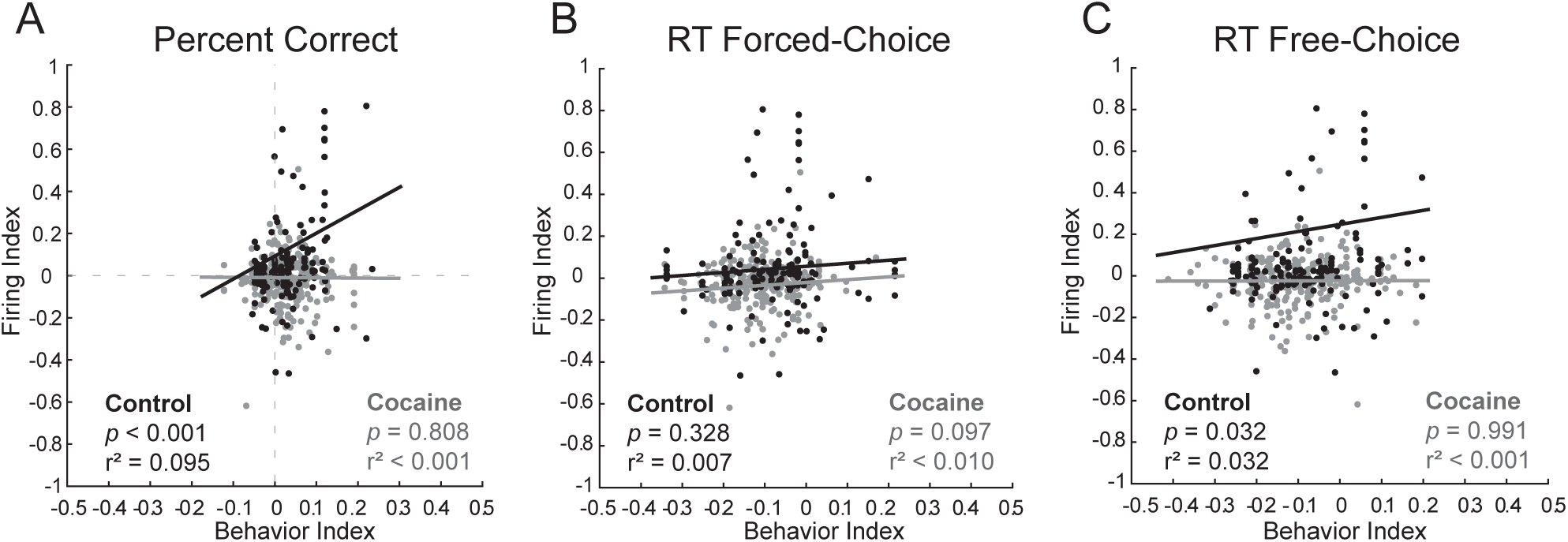
Several relationships between behavior and size-encoding activity are disrupted by cocaine exposure. Scatter plots of context index (size block – delay block/ size block + delay block) distribution data from neurons that increased activity during the reward epoch compared to ratios of size vs delay behavior for **A**. Percent Correct, **B**. Forced-choice reaction times, and **C**. Free-choice reaction times. Black = control; Gray = Cocaine.

As for reaction time, we observed no significant correlations between behavior and firing during forced-choice trials for either controls (*Regression, p* = 0.328, *r*^2^ = 0.007) or cocaine-exposed rats (*Regression, p* = 0.097; *Fig. 6B*). However, during free-choice trials there was a significant, positive relationship between firing rate and reaction time in control rats (*Regression, p* = 0.032, *r*^2^ = 0.032). This result suggests that stronger firing rates were actually related to slower reaction times during size blocks. This relationship was disrupted by cocaine exposure (*Regression, p* = 0.991, *r*^2^ < 0.001; *Fig. 6C*), and was significantly different from what was observed in controls (*Fisher r-to-z transformation*; z = 1.72, *p* = 0.043).

## DISCUSSION

Insula’s traditional roles integrating experiences like auditory perception (Kimura et al., 2010), gustation (Hanamori et al., 1998), and pain (Downar et al., 2003) with internal state have expanded to reward-guided decision-making. Previous recording studies, for example, demonstrate insula activity encodes anticipation of both positive (Samuelsen et al., 2013, 2012) and negative consequences (Jo and Jung, 2016), and reward consumption (Kusumoto-Yoshida et al., 2015; Wittmann et al., 2020). In relation to its interoceptive functions, insula also seems to track the subjective value of a reward by reflecting how context and experience can change a reward’s value even when the reward itself has not changed. Satiety, for example, reduces insula activity for both food (Livneh et al., 2017) and water (Livneh et al., 2020; Namboodiri and Stuber, 2020). Other work has shown that when a previously appetitive sucrose solution is paired with delayed (as opposed to immediate) access to cocaine, insula activity also seems to reflect the reduced preference of this once favorable reward, even though the sucrose itself has not changed (Moschak et al., 2018). Our work expands on these studies by demonstrating signals in the insula related to cues, context, and differently valued rewards, and how chronic cocaine exposure profoundly affects insula activity in reward-guided decision-making. First, we see enhanced signals representing reward, especially processing related to the delivery of immediate reward. Second, cocaine exposure reduces signals related to cue and block context selectivity, suggesting insula plays an important part in drug-induced impairments of both executive control and decision-making.

Similar to recent work from the Fontanini group (Vincis et al., 2020), our study found that insula encoded predicted outcomes during the presentation of reward-predictive odor cues, and fired in anticipation of reward after completion of an instrumental response. Further, we showed that firing was higher for cues that predicted high-valued reward in both delay and size domains. These neurons likely contributed to subjective preferences of more immediate reward and the maintenance of reward representations across long delays. Interestingly, these neural patterns of firing are similar to what we have reported previously in OFC and NAc (Burton et al., 2018b, 2014c; Takahashi et al., 2009). Both these regions are outcome selective during cues, and fire during the anticipation and delivery of reward. However, unique to insula, firing did not differ between rewards delivered after short and long delays in control rats. Thus, in normal insula, anticipatory reward signals were not discounted by the length of the delay that preceded it in our task.

Another striking characteristic unique to insula is that average firing was stronger during size blocks, and correlated with changes in reaction time and accuracy. When designing the behavioral task, it was our goal to completely counter-balance value manipulation and direction within each session. We could only accomplish this objective by running size blocks last, countering satiation effects by promising rats more and faster access to reward later in each behavioral session. Indeed, rats actually exhibited faster reaction times and better percent correct scores during size blocks due to the increased benefit of completing trials (i.e., no long delays and potential for a larger reward). However, higher firing during size blocks reflects more than just receiving a larger reward within trials, as activity was stronger before initiation of trials and after delivery of reward on ‘small-reward’ trials, which were physically identical to reward delivered on ‘short-delay’ trials (i.e., both trial types delivered 1 bolus after 500ms delay). We hypothesize that these increases in activity track the overall value of block contexts and contribute to elevating motivation as satiation sets in, based on insula activity’s correlations to changes in both reaction time and percent correct. Such interpretations are in line with a recent publication from Wittman et al. (2020), which also implicates insula in tracking long-term reward; their task demonstrates a relationship between BOLD signaling in the macaque insula and task engagement during blocks of trials where probability of reward receipt is overall either higher or lower. Our context effects suggest such global reward signals in insula are encoded at the single-cell level, and may also represent specific reward outcome in addition to reward probability. Moreover, our work adds to a small but growing pool of physiological evidence supporting long-held theories that insula contributes to interoception by connecting external stimuli and internal experience, and by making context-based adjustments to representations of reward value in order to optimize behavior (Droutman et al., 2015; Naqvi and Bechara, 2010).

In addition to reporting these novel aspects of insula encoding in control rats, we also determined how these correlates are disrupted after chronic cocaine self-administration. Our results replicate previous behavioral findings demonstrating that rats in both cocaine and control groups prefer high-valued rewards, and that this preference is exaggerated in cocaine-exposed animals (Burton et al., 2018b, 2017; Roesch et al., 2007; Roesch and Bryden, 2011; Vázquez et al., 2019). Interestingly, prior cocaine self-administration shifted the balance of encoding in insula from cues and context to reward. That is, in rats that had self-administered cocaine, there were fewer neurons that increased firing to cues and context. Instead, after cocaine exposure, significantly more neurons increased firing in anticipation of more immediate reward. This activity pattern might encourage impulsive choice by not adequately conveying the value of differently delayed and sized rewards at the time of the decision and more strongly represent the anticipation that reward should have been delivered at the shorter delay. Consistent with this hypothesis, not only did insula neurons fire more strongly in anticipation of reward on short delay trials, but they also fired strongly at the start of long delay trials at the time when reward would have been delivered on trials with shorter delays (i.e., 500 ms after well entry).

Insula neurons in cocaine-exposed rats also differed from controls in that they failed to increase firing at the end of recording sessions during size trial blocks. Above, we suggest that this signal is important for motivating satiated animals when they encounter global increases in reward value in the context of size blocks. Remarkably, this signal was not present in the insula of rats that had self-administered cocaine. The explanation for this finding may be that these signals (i.e., signals related to satiety, motivation or absolute value) necessary to drive behavior during this phase in the task are disrupted after cocaine exposure. Another interpretation may be that these size-encoding signals are absent because they are not necessary to drive motivation in cocaine-exposed rats, as seen by their faster reaction times and higher accuracy in size blocks. However, the absence of these global signals may in turn disrupt the processing of long vs short-delay rewards and subsequently bias behavior for immediate rewards. This proposed reduction of context and task structure representations in favor of reward fits in well with theories of drug exposure enhancing model-free behavior (Dayan and Berridge, 2011; Everitt and Robbins, 2005; Jentsch and Taylor, 1999; Lucantonio et al., 2014; Volkow and Fowler, 2000), and could be explored further by examining insula activity in other behavioral tasks, including those in which the selection of immediate reward is not always the most favorable action.

To conclude, our findings demonstrate that cocaine exposure enhances signals related to reward delivery while attenuating signals related to cue and context. Our novel context encoding results-insula reflecting changes in global reward value as animals become satiated-provide physiological evidence of insula’s role in maintaining internal states in relation to external environmental changes (Naqvi and Bechara, 2010). Moreover, the disruptions of these signals and related changes in behavior due to cocaine exposure suggest an important role for insula in impaired decision-making observed after drug abuse. Further study of insula using different behavioral assays will be important for understanding how this region is involved in optimal decision-making, and its drug-induced impairments (Dayan and Berridge, 2011; Everitt and Robbins, 2005; Jentsch and Taylor, 1999; Lucantonio et al., 2014; Volkow and Fowler, 2000).

## MATERIALS AND METHODS

### Subjects

Nine Long-Evans rats (weight: 175-200g, 3 females; 6 males) rats were obtained from Charles River Laboratories. Subjects were held and tested at University of Maryland, College Park according to university and NIH guidelines.

### Odor-guided delay/size choice task

Rats were trained on the Delay/Size task (summarized in *Fig. 1*) for one month prior to surgery. In brief, subjects needed to nose poke into a central port during house light illumination to receive one of three odor cues (odors were 2-Octanol, Pentyl Acetate, or Carvone). These odors instruct rats which direction to move to receive a 10% liquid sucrose reward from wells located on either side of the central port. Two of these odors instructed rats to go either left or right for reward (forced-choice), while the third odor indicated that rats could receive a reward from either well (free-choice). Forced-choice odors were counterbalanced across rats, and were presented in a pseudorandom sequence with free-choice odor presented on 7/20 trials. Incorrect well-selection during a forced-choice trials resulted in no reward delivery, and 500ms time-out where the subject could not nose poke and start a new trial.

During each recording session, reward value was independently manipulated across four blocks of 60 correct trials (Trial summary in *Fig. 1A*; Block summary in *Fig. 1B*). During the first two blocks, one well was randomly designated to deliver reward immediately (500ms delay; 0.05ml)) while the other well delivered sucrose with delays that would gradually increase (1000-7000ms). Delay-well contingencies would then be switched at the start of the second block so that the previously higher-valued well carried the longer delay. During the final two trial blocks, delays on both wells were held constant at 500ms, and reward value was instead manipulated by size. At the start of the third block, the well that previously carried the long delay condition now delivered two boli of sucrose solution separated by 500 ms. Value contingencies were again switched at the fourth block (i.e, large became small and small became large). Subjects were water-deprived to encourage motivation during the task.

### Surgery

All subjects were implanted with catheters in the jugular vein for self-administration and drivable electrodes (+1.5AP, +/-5.0ML, −5.0DV from brain surface) for single-unit recordings (control = 5 and cocaine = 4). Catheters were made from SILASTIC tubing (0.02×0.037 in; Dow Corning, Midland, MI) with a modified G 5-up cannula (Plastics One, Roanoke, VA). The cannula traveled through the fascia layer over the shoulder and was cemented on top of the skull. Electrodes (bundles of 10-25µm-diameter FeNiCr wire) were implanted during this same surgery.

### Self-Administration

After one week of recovery, rats underwent a 12 d self-administration protocol in Med Associates operant chambers (St. Albans, VT). Animals in the cocaine group could press a lever for an infusion of cocaine, while control animals pressed for sucrose pellets. On days 1-6, rats self-administered 1 mg/kg dosage of cocaine or 2 sucrose pellets per lever press for a maximum of 30 infusions or for a 3 h time limit. For the final 6 days, the dosage of cocaine was halved to 0.5mg/kg and only 1 sucrose pellet was delivered per lever press. This procedure allowed us to assess increases in drug-seeking when doses are cut in half to maintain the desired level of drug intake. All self-administration procedures are consistent with work establishing that continuous access to high cocaine doses evokes drug-taking and drug-seeking behaviors that are consistent with promoting symptoms of addiction (Allain and Samaha, 2019), and has been shown to change behavior and neural signals in other brain regions (Burton et al., 2018b, 2017; Vázquez et al., 2019).

### Single-unit recording

Procedures were the same as described previously (Bryden and Roesch, 2015). Wires were screened for activity each day; if no activity was detected, the rat was removed and the electrode assembly was advanced 40 or 80µm. Otherwise, a session was conducted, and the electrode was advanced at the end of the session. Neural activity was recorded using four identical Plexon Multichannel Acquisition Processor systems (Dallas, TX). Signals from electrode wires were amplified 20x by an op-amp headstage, located on the electrode array. Immediately outside the training chamber, the signals were passed through a differential pre-amplifier (Plexon Inc, PBX2/16sp-r-G50/16fp-G50) where single unit signals were amplified 50x and filtered at 150-9000Hz. The single unit signals were then sent to the Multichannel Acquisition Processor box, where they were further filtered at 250-8000Hz, digitized at 40kHz and amplified at 1-32x. Waveforms (>2.5:1 signal-to-noise) were extracted from active channels and recorded to the disk by an associated workstation with event timestamps *from the behavior computer*.

### Behavioral Analysis

Behavior in the recording task was analyzed by calculating the percent of correct responses on forced-choice trials (i.e. the amount of trials the animal correctly responded to the side corresponding to the directional odor cue), the percent of trials rats chose a particular valued condition (short, long, large, small) on free-choice trials, and reaction times (odor offset to odor port exit). Behavioral analyses were computed for each individual session (separated by cocaine and control groups), and then averaged across sessions for each group. Conducting analyses across sessions—instead of across individual subjects—provides a better reflection of the neural correlates corresponding to behavior. Importantly, the main behavioral findings described in this paper have been replicated in three different studies (Burton et al., 2018b, 2017; Vázquez et al., 2019). Multi-factor analysis of variance factors included group (control vs. cocaine), reward value (high vs. low), and value manipulation (size vs. delay). Post hoc t-tests (p<0.05) were used to determine differences between the cocaine and control groups.

### Neural Analysis

Single units were analyzed in Offline Sorter using template matching software (Plexon), and were exported to NeuroExplorer (Nex Technologies, Colorado Springs,CO) and Matlab (MathWorks, Natick, MA). Only rewarded trials were analyzed, from both free and forced-choice trials. Neural activity was analyzed in three epochs: baseline firing rate, odor cue epoch, and reward epoch. All were calculated by dividing the total number of spikes by time. Baseline activity was taken 1 s before odor presentation, the odor epoch was taken 100ms after odor onset until port exit, and the reward epoch was taken 250ms before sucrose delivery to 1s after reward delivery. The reward epoch was designed to capture activity related to reward expectancy and delivery. There was no overlap between epochs, even during short delay (500ms) conditions. To analyze neural activity during these epochs we normalized firing rates so that high and low-firing cells could be analyzed together. We determined significant changes in firing rate by taking difference scores for each neuron, and plotting the number of neurons with significant differential firing from baseline (demonstrated through Wilcoxon tests, *p* < 0.05, black bars in distributions). Relationships between neural firing and behavioral activity were determined using regression analyses.

Upon completion of recordings, subjects were euthanized via isoflurane overdose and perfused transcardially with 200ml of 0.9% saline, followed by 500ml of 4% paraformaldehyde (PFA) in order to fix brain tissue and confirm electrode placement (*Fig. 1D*).

## ACKNOWLEDGEMNTS

This work was supported by the National Institutes of Health National Institute on Drug Abuse Grant R01 DA031695 to Dr. Matthew Roesch.

## REFERENCES

Allain F, Samaha AN. 2019. Revisiting long-access versus short-access cocaine self-administration in rats: intermittent intake promotes addiction symptoms independent of session length. Addict Biol 24:641–651. doi:10.1111/adb.12629

Bonson KR, Grant SJ, Contoreggi CS, Links JM, Metcalfe J, Weyl HL, Kurian V, Ernst M, London ED. 2002. Neural Systems and Cue-Induced Cocaine Craving. Neuropsychopharmacology 26:376–386.

Brockett AT, Pribut HJ, Vázquez D, Roesch MR. 2018. The impact of drugs of abuse on executive function: characterizing long-term changes in neural correlates following chronic drug exposure and withdrawal in rats. Learn Mem 25:461–473. doi:10.1101/lm.047001.117

Bryden DW, Johnson EE, Diao X, Roesch MR. 2011. Impact of expected value on neural activity in rat substantia nigra pars reticulata. Eur J Neurosci 33:2308–2317. doi:10.1111/j.1460-9568.2011.07705.x

Bryden DW, Roesch MR. 2015. Executive Control Signals in Orbitofrontal Cortex during Response Inhibition. J Neurosci 35:3903–3914. doi:10.1523/JNEUROSCI.3587-14.2015

Burke CJ, Tobler PN. 2011. Reward skewness coding in the insula independent of probability and loss. J Neurophysiol 106:2415–22. doi:10.1152/jn.00471.2011

Burton AC, Bissonette GB, Lichtenberg NT, Kashtelyan V, Roesch MR. 2014a. Ventral striatum lesions enhance stimulus and response encoding in dorsal striatum. Biol Psychiatry 75:132–139. doi:10.1016/j.biopsych.2013.05.023

Burton AC, Bissonette GB, Vazquez D, Blume EM, Donnelly M, Heatley KC, Hinduja A, Roesch MR. 2018a. Previous cocaine self-administration disrupts reward expectancy encoding in ventral striatum. Neuropsychopharmacology 43:2350–2360. doi:10.1038/s41386-018-0058-0

Burton AC, Bissonette GB, Vazquez D, Blume EM, Donnelly M, Heatley KC, Hinduja A, Roesch MR. 2018b. Previous cocaine self-administration disrupts reward expectancy encoding in ventral striatum. Neuropsychopharmacology 43:2350–2360. doi:10.1038/s41386-018-0058-0

Burton AC, Bissonette GB, Zhao AC, Patel PK, Roesch MR. 2017. Prior cocaine self-administration increases response–outcome encoding that is divorced from actions selected in dorsal lateral striatum. J Neurosci 37:7737–7747. doi:10.1523/JNEUROSCI.0897-17.2017

Burton AC, Kashtelyan V, Bryden DW, Roesch MR. 2014b. Increased firing to cues that predict low-value reward in the medial orbitofrontal cortex. Cereb Cortex 24:3310–3321. doi:10.1093/cercor/bht189

Burton AC, Kashtelyan V, Bryden DW, Roesch MR. 2014c. Increased firing to cues that predict low-value reward in the medial orbitofrontal cortex. Cereb Cortex 24:3310–3321. doi:10.1093/cercor/bht189

Burton AC, Nakamura K, Roesch MR. 2015. From ventral-medial to dorsal-lateral striatum: neural correlates of reward-guided decision-making. Neurobiol Learn Mem 117:51–9. doi:10.1016/j.nlm.2014.05.003

Campbell EJ, Flanagan JPM, Walker LC, Hill MKRI, Marchant NJ, Lawrence AJ. 2019. Anterior insular cortex is critical for the propensity to relapse following punishment-imposed abstinence of alcohol seeking. J Neurosci 39:1077–1087. doi:10.1523/JNEUROSCI.1596-18.2018

Cofresí RU, Grote DJ, Viet Thanh Le E, Monfils M-H, Chaudhri N, Gonzales RA, Lee HJ. 2018. Alcohol-associated antecedent stimuli elicit alcohol seeking in non-dependent rats and may activate the insula. Alcohol. doi:10.1016/J.ALCOHOL.2018.08.004

Contreras M, Ceric F, Torrealba F. 2007. Inactivation of the Interoceptive Insula Disrupts Drug Craving and Malaise Induced by Lithium. Proc Natl Acad Sci USA 104. doi:10.1126/science.1147248

Craig AD. 2010. The sentient self. Brain Struct Funct 214:563–577. doi:10.1007/s00429-010-0248-y

Craig AD. 2002. How do you feel? Interoception: the sense of the physiological condition of the body. Nat Rev Neurosci 3:655–666. doi:10.1038/nrn894

Damasio AR, Grabowski TJ, Bechara A, Damasio H, Ponto LLB, Parvizi J, Hichwa RD. 2000. Subcortical and cortical brain activity during the feeling of self-generated emotions. Nat Neurosci 3:1049–1056. doi:10.1038/79871

Dayan P, Berridge KC. 2011. Model-based and model-free Pavlovian reward learning: Revaluation, revision, and revelation. Cogn Affect Behav Neurosci 14:473–492. doi:10.3758/s13415-014-0277-8

Downar J, Mikulis DJ, Davis KD. 2003. Neural correlates of the prolonged salience of painful stimulation. Neuroimage 20:1540–1551. doi:10.1016/S1053-8119(03)00407-5

Droutman V, Read SJ, Bechara A. 2015. Revisiting the role of the insula in addiction. Trends Cogn Sci 19:414–420. doi:10.1016/j.tics.2015.05.005

Everitt BJ, Robbins TW. 2005. Neural systems of reinforcement for drug addiction: from actions to habits to compulsion. Nat Neurosci 8:1481–1489. doi:10.1038/nn1579

Forget B, Pushparaj A, Le Foll B. 2010. Granular Insular Cortex Inactivation as a Novel Therapeutic Strategy for Nicotine Addiction. Biol Psychiatry 68:265–271. doi:10.1016/j.biopsych.2010.01.029

García R, Simón MJ, Puerto A. 2013. Conditioned place preference induced by electrical stimulation of the insular cortex: effects of naloxone. Exp Brain Res 226:165–174. doi:10.1007/s00221-013-3422-7

Gil-Lievana E, Balderas I, Moreno-Castilla P, Luis-Islas J, McDevitt RA, Tecuapetla F, Gutierrez R, Bonci A, Bermúdez-Rattoni F. 2020. Glutamatergic basolateral amygdala to anterior insular cortex circuitry maintains rewarding contextual memory. Commun Biol 3:1–11. doi:10.1038/s42003-020-0862-z

Guillem K, Kravitz A V., Moorman DE, Peoples LL. 2010. Orbitofrontal and insular cortex: Neural responses to cocaine-associated cues and cocaine self-administration. Synapse 64:1–13. doi:10.1002/syn.20698

Hanamori T, Kunitake T, Kato K, Kannan H. 1998. Neurons in the posterior insular cortex are responsive to gustatory stimulation of the pharyngolarynx, baroreceptor and chemoreceptor stimulation, and tail pinch in rats. Brain Res 785:97–106. doi:10.1016/S0006-8993(97)01392-9

Jentsch JD, Taylor JR. 1999. Impulsivity resulting from frontostriatal dysfunction in drug abuse: implications for the control of behavior by reward-related stimuli. Psychopharmacology (Berl) 146:373–390. doi:10.1007/PL00005483

Jo S, Jung MW. 2016. Differential coding of uncertain reward in rat insular and orbitofrontal cortex. Sci Rep 6:1–13. doi:10.1038/srep24085

Kashtelyan V, Tobia SC, Burton AC, Bryden DW, Roesch MR. 2012. Basolateral amygdala encodes upcoming errors but not response conflict. Eur J Neurosci 35:952–9. doi:10.1111/j.1460-9568.2012.08022.x

Kimura A, Imbe H, Donishi T. 2010. Efferent connections of an auditory area in the caudal insular cortex of the rat: anatomical nodes for cortical streams of auditory processing and cross-modal sensory interactions. Neuroscience 166:1140–1157. doi:10.1016/j.neuroscience.2010.01.032

Kusumoto-Yoshida I, Liu H, Chen BT, Fontanini A, Bonci A. 2015. Central role for the insular cortex in mediating conditioned responses to anticipatory cues. PNAS 112:1190–1195. doi:10.1073/pnas.1416573112

Li CL, Zhu N, Meng XL, Li YH, Sui N. 2013. Effects of inactivating the agranular or granular insular cortex on the acquisition of the morphine-induced conditioned place preference and naloxone-precipitated conditioned place aversion in rats. J Psychopharmacol 27:837–844. doi:10.1177/0269881113492028

Livneh Y, Ramesh RN, Burgess christian R, Levandowski KM, Madara JC, Fenselau H, Goldey GJ, Diaz VE, Jikomes N, Resch JM, Lowell BB, Andermann ML. 2017. Homeostatic circuits selectively gate food cue responses in insular cortex. Nature 546:611–616. doi:10.1038/nature22375

Livneh Y, Sugden AU, Madara JC, Essner RA, Flores VI, Sugden LA, Resch JM, Lowell BB, Andermann ML. 2020. Estimation of Current and Future Physiological States in Insular Cortex. Neuron 105:1094-1111.e10. doi:10.1016/j.neuron.2019.12.027

Lucantonio F, Caprioli D, Schoenbaum G. 2014. Transition from ‘model-based’ to ‘model-free’ behavioral control in addiction: Involvement of the orbitofrontal cortex and dorsolateral striatum. Neuropharmacology 76:407–415. doi:10.1016/j.neuropharm.2013.05.033

Mendez IA, Simon NW, Hart N, Mitchell MR, Nation JR, Wellman PJ, Setlow B. 2010. Self-administered cocaine causes long-lasting increases in impulsive choice in a delay discounting task. Behav Neurosci 124:470–477. doi:10.1037/a0020458

Mesulam M-M, Mufson EJ. 1982. Insula of the old world monkey. III: Efferent cortical output and comments on function. J Comp Neurol 212:38–52. doi:10.1002/cne.902120104

Mizoguchi H, Katahira K, Inutsuka A, Fukumoto K, Nakamura A, Wang T, Nagai T, Sato J, Sawada M, Ohira H, Yamanaka A, Yamada K. 2015. Insular neural system controls decision-making in healthy and methamphetamine-treated rats. Proc Natl Acad Sci U S A 112:E3930–E3939. doi:10.1073/pnas.1418014112

Moschak TM, Wang X, Carelli RM. 2018. A neuronal ensemble in the rostral agranular insula tracks cocaine-induced devaluation of natural reward and predicts cocaine seeking. J Neurosci 38:8463–8472. doi:10.1523/JNEUROSCI.1195-18.2018

Namboodiri VMK, Stuber GD. 2020. Interoceptive Inception in Insula. Neuron 105:959–960. doi:10.1016/j.neuron.2020.02.032

Naqvi NH, Bechara A. 2010. The insula and drug addiction: an interoceptive view of pleasure, urges, and decision-making. Brain Struct Funct 214:435–450. doi:10.1007/s00429-010-0268-7

Naqvi NH, Rudrauf D, Damasio H, Bechara A. 2007. Damage to the insula disrupts addiction to cigarette smoking. Science (80-) 315:531–534. doi:10.1126/science.1135926

Ongur D, Price JL. 2000. The Organization of Networks within the Orbital and Medial Prefrontal Cortex of Rats, Monkeys and Humans. Cereb Cortex 10:206–219. doi:10.1093/cercor/10.3.206

Pelloux Y, Murray JE, Everitt BJ. 2013. Differential roles of the prefrontal cortical subregions and basolateral amygdala in compulsive cocaine seeking and relapse after voluntary abstinence in rats. Eur J Neurosci 38:3018–3026. doi:10.1111/ejn.12289

Perdomo G, Contreras M, Torrealba F, Billeke P, Madrid C, Vicencio S, González M. 2012. A Role for the Insular Cortex in Long-Term Memory for Context-Evoked Drug Craving in Rats. Neuropsychopharmacology 37:2101–2108. doi:10.1038/npp.2012.59

Preuschoff K, Quartz SR, Bossaerts P. 2008. Human Insula Activation Reflects Risk Prediction Errors As Well As Risk. doi:10.1523/JNEUROSCI.4286-07.2008

Pushparaj A, Le Foll B. 2015. Involvement of the caudal granular insular cortex in alcohol self-administration in rats. Behav Brain Res 293:203–207. doi:10.1016/J.BBR.2015.07.044

Roesch MR, Bryden DW. 2011. Impact of size and delay on neural activity in the rat limbic corticostriatal system. Front Neurosci 5:1–13. doi:10.3389/fnins.2011.00130

Roesch MR, Bryden DW, Cerri DH, Haney ZR, Schoenbaum G. 2012. Willingness to wait and altered encoding of time-discounted reward in the orbitofrontal cortex with normal aging. J Neurosci 32:5525–5533. doi:10.1523/JNEUROSCI.0586-12.2012

Roesch MR, Takahashi Y, Gugsa N, Bissonette GB, Schoenbaum G. 2007. Previous cocaine exposure makes rats hypersensitive to both delay and reward magnitude. J Neurosci 27:245–250. doi:10.1523/JNEUROSCI.4080-06.2007

Roesch MR, Taylor AR, Schoenbaum G. 2006. Encoding of Time-Discounted Rewards in Orbitofrontal Cortex Is Independent of Value Representation. Neuron 51:509–520. doi:10.1016/j.neuron.2006.06.027

Rogers-Carter MM, Christianson JP. 2019. An Insular View of the Social Decision-Making Network. Neurosci Biobehav Rev. doi:10.1016/J.NEUBIOREV.2019.06.005

Samuelsen CL, Gardner MPH, Fontanini A. 2013. Thalamic contribution to cortical processing of taste and expectation. J Neurosci 33:1815–1827. doi:10.1523/JNEUROSCI.4026-12.2013

Samuelsen CL, Gardner MPH, Fontanini A. 2012. Effects of Cue-Triggered Expectation on Cortical Processing of Taste. Neuron 74:410–422. doi:10.1016/j.neuron.2012.02.031

Seif T, Chang S-J, Simms JA, Gibb SL, Dadgar J, Chen BT, Harvey BK, Ron D, Messing RO, Bonci A, Hopf FW. 2013. Cortical activation of accumbens hyperpolarization-active NMDARs mediates aversion-resistant alcohol intake. Nat Neurosci 16:1094–1100. doi:10.1038/nn.3445

Simon NW, Mendez IA, Setlow B. 2007. Cocaine exposure causes long-term increases in impulsive choice. Behav Neurosci 121:543–549. doi:10.1037/0735-7044.121.3.543

Takahashi YK, Roesch MR, Stalnaker TA, Haney RZ, Calu DJ, Taylor AR, Burke KA, Schoenbaum G. 2009. The Orbitofrontal Cortex and Ventral Tegmental Area Are Necessary for Learning from Unexpected Outcomes. Neuron 62:269–280. doi:10.1016/j.neuron.2009.03.005

Takahashi YK, Stalnaker TA, Roesch MR, Schoenbaum G. 2017. Effects of inference on dopaminergic prediction errors depend on orbitofrontal processing. Behav Neurosci 131:127–134. doi:10.1037/bne0000192

Vázquez D, Pribut HJ, Burton AC, Tennyson SS, Roesch MR. 2019. Prior cocaine self-administration impairs attention signals in anterior cingulate cortex. Neuropsychopharmacology. doi:10.1038/s41386-019-0578-2

Vincis R, Chen K, Czarnecki L, Chen J, Fontanini A. 2020. Dynamic Representation of Taste-Related Decisions in the Gustatory Insular Cortex of Mice. Curr Biol 30:1834-1844.e5. doi:10.1016/j.cub.2020.03.012

Volkow ND, Fowler JS. 2000. Addiction, a Disease of Compulsion and Drive: Involvement of the Orbitofrontal Cortex. Cereb Cortex 10:318–325. doi:10.1093/cercor/10.3.318

Wang G, Volkow N, Fowler J, Cervany P, Hitzemann R, Pappas N, Wong C, Felder C. 1999. Regional brain metabolic activation during craving elicited by recall of previous drug experiences, Life Sciences. Elsevier Science Inc.

Wittmann MK, Fouragnan E, Folloni D, Klein-flügge MC, Chau BKH, Khamassi M, Rushworth MFS. 2020. Global reward state affects learning and activity in raphe nucleus and anterior insula in monkeys. Nat Commun 11:1–17. doi:10.1038/s41467-020-17343-w

Zhang R, Jia W, Wang Y, Zhu Y, Liu F, Li B, Liu F, Wang H, Tan Q. 2019. A glutamatergic insular-striatal projection regulates the reinstatement of cue-associated morphine-seeking behavior in mice. Brain Res Bull 152:257–264. doi:10.1016/j.brainresbull.2019.07.023

